# Empirical prediction of variant-associated cryptic-donors with 87% sensitivity and 95% specificity

**DOI:** 10.1101/2021.07.18.452855

**Authors:** Ruebena Dawes, Himanshu Joshi, Sandra T. Cooper

## Abstract

Predicting which cryptic-donors may be activated by a genetic variant is notoriously difficult. Through analysis of 5,145 cryptic-donors activated by 4,811 variants (versus 86,963 decoy-donors not used; any GT or GC), we define an empirical method predicting cryptic-donor activation with 87% sensitivity and 95% specificity. Strength (according to four algorithms) and proximity to the authentic-donor appear important determinants of cryptic-donor activation. However, other factors such as auxiliary splicing elements, which are difficult to identify, play an important role and are likely responsible for current prediction inaccuracies. We find that the most frequent mis-splicing events at each exon-intron junction, mined from 40,233 RNA-sequencing samples, predict with remarkable accuracy which cryptic-donor will be activated in rare disease. Aggregate RNA-Sequencing splice-junction data provides an accurate, evidence-based method to predict variant-activated cryptic-donors in genetic disorders, assisting pathology consideration of possible consequences of a variant for the encoded protein and RNA diagnostic testing strategies.

## Introduction

Genetic variants affecting the conserved sequences of the consensus splicing motifs can alter binding of spliceosomal components and induce mis-splicing of precursor messenger RNA (pre-mRNA)^1^, making them a common cause of inherited disorders^2–5^. Splicing variants can simultaneously induce different mis-splicing outcomes, including skipping of one or more exons, activation of a cryptic splice-site(s), and/or retention of one or more introns^1^. Whether induced mis-splicing disrupts the reading frame or affects a region of known functional (and clinical) importance, has significant diagnostic implications. Therefore, knowing the specific mis-splicing outcome of genetic variant is necessary to conclusively link it to a disease. While the accuracy of *in silico* algorithms in predicting whether a variant will cause mis-splicing has been comprehensively assessed^6–9^, there is currently no reliable means to predict *which* mis-splicing event(s) may occur in response to a variant that activates mis-splicing. As a result of this and other factors, the vast majority of splice site variants are classified as variants of uncertain significance (VUS); a non-actionable diagnostic endpoint in genomic medicine^10^.

We recently evaluated the accuracy and concordance of SpliceAI (SAI)^11^ and algorithms within Alamut Visual^®^ (Interactive Biosoftware, Rouen, France)^12,13^, to predict splicing outcomes arising from genetic variants identified in 74 families with monogenetic conditions subject to RNA diagnostic studies (79 variants; 19% essential GT-AG splice-site variants and 71% extended splice-site variants)^14^. Algorithmic predictions of the strengths of activated cryptic splice sites were highly discordant, especially for cryptic-donors. SAI’s deep learning showed the greatest accuracy in predicting activated cryptic splice-site(s) (66% true positive with 34% false negative), whereas historical algorithms within Alamut Visual^®^ resulted in 34% - 69% false negatives^14^.

In this study we focus on determining empirical features that inform prediction of variant-associated spliceosomal selection of a cryptic-donor, in preference to the ‘authentic-donor’ (positioned at the exon-intron junction), and other nearby decoy-donors (any GT or GC) that are not used by the spliceosome. Through analysis of 4,811 variants in 3,399 genes, we show that while intrinsic splice-site strength and proximity to the authentic-donor strongly influence spliceosomal selection of a cryptic-donor, these factors alone are not sufficient for accurate prediction. Importantly, we show that the most frequent mis-splicing events seen at each exon-intron junction across 40,233 publicly available RNA-seq samples predict with remarkable accuracy which cryptic-donor will be activated in rare disease. An improved ability to reliably identify cryptic-donors will greatly assist both theoretical consideration of possible consequences for the encoded protein, as well as guide PCR-based RNA-diagnostic studies seeking to experimentally determine consequences for pre-mRNA splicing^14^. In turn this will improve our ability to classify splicing variants, avoiding the non-actionable diagnostic endpoint of a variant of uncertain significance (VUS).

## Results

### Reference database of variants activating a cryptic-donor

We collate a database of cryptic-donor variants, defined as variant-associated erroneous use of a donor other than the authentic-donor (positioned at the exon-intron junction). Variants were derived from several sources^11,16,17^ (Figure 1A, upper. See methods). The genomic locations and extended donor consensus sequences of the authentic-donor, cryptic-donor(s), as well as any decoy-donors (any GT/GC motif) within 250 nucleotides (nt) of authentic-donors, were compiled for analysis. We define the extended donor splice-site region as spanning the fourth to last nucleotide of the exon (E^−4^, E = exon) to the eighth nucleotide of the intron (D^+8^; D = donor) (see Figure 1A).

**Fig 1.**
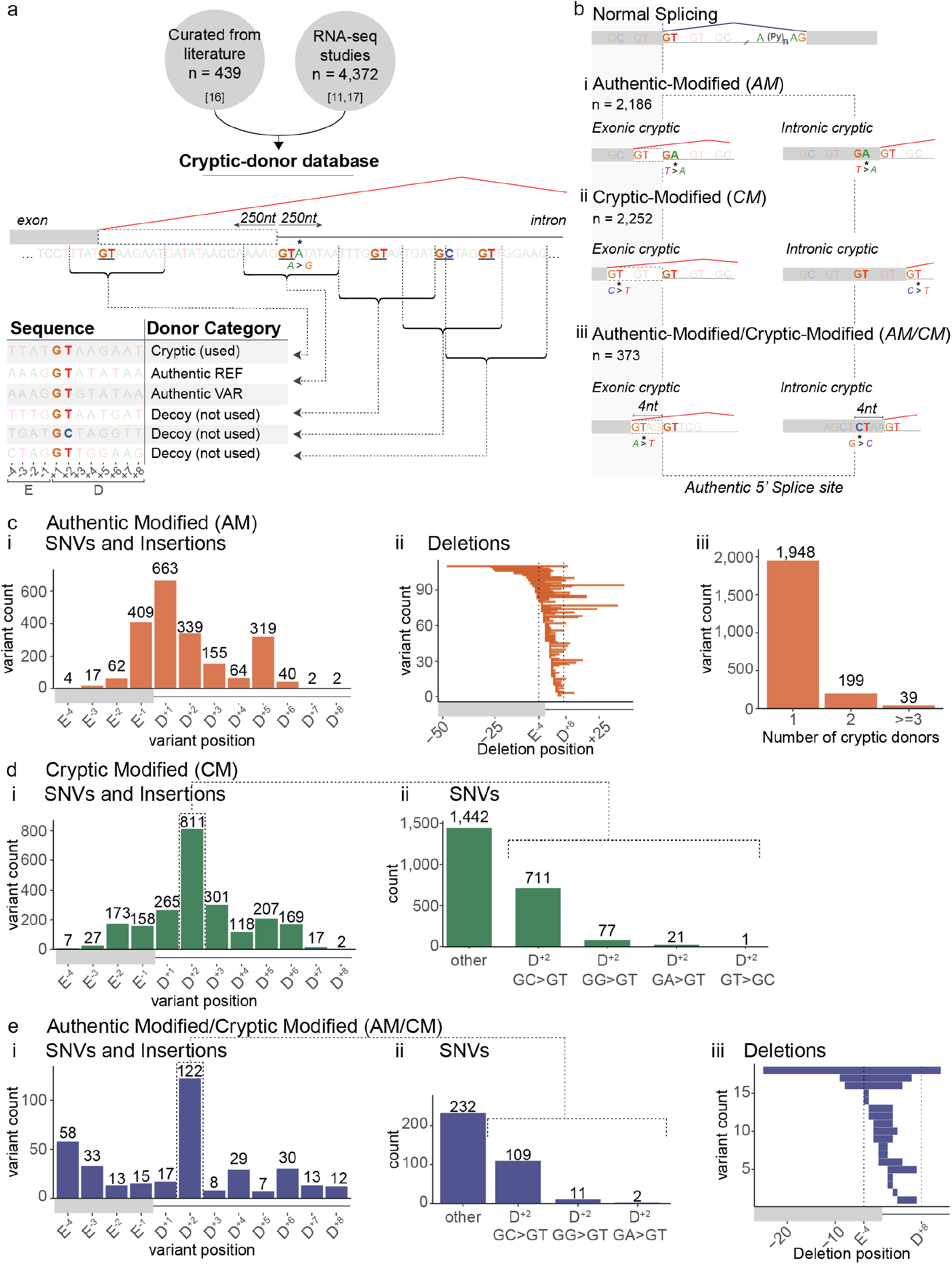
Reference database of variants activating a cryptic-donor. **a)** Schematic of the cryptic-donor database. See methods for details. **b)** Schematic of the three categories of cryptic-donor variants in the database: ***i)*** Authentic Modified (*AM*-variants), ***ii)*** Cryptic Modified (*CM*-variants) and ***iii)*** Authentic Modified and Cryptic Modified (*AM/CM*-variants). **c)** Characteristics of *AM*-variants (n = 2,186, *orange*). Positions of ***(i)*** Single Nucleotide Variants (SNVs), insertions and ***(ii)*** deletions relative to the authentic-donor splice-site. In ***(ii)*** each of the horizontal bars represents one deletion variant showing the position and width of each deletion relative to the authentic-donor. ***iii)*** The number of cryptic-donors activated by each *AM*-variant. **d)** Characteristics of *CM*-variants (n = 2,255; green). ***i)*** Positions of SNVs and insertions relative to the cryptic-donor. ***ii)*** Frequency of SNVs resulting in cryptic activation, highlighting the prevalence or GC>GT D^+2^ variants (i.e. the +2 position of the donor splice-site). **e)** Characteristics of *AM/CM*-variants (n = 373, *blue*). ***i-ii)*** as in **d*i-ii. iii)*** as in **c*ii***.

Cryptic-donor variants fall into three categories (Figure 1B): ***A)*** *‘Authentic-Modified’ (AM)*: a genetic variant modifies the authentic-donor resulting in activation one or more cryptic-donors already present in the genome (n = 2,186) (Figure 1Bi). *AM*-variants commonly affect the E^−1^, D^+1^, D^+2^ and D^+5^ positions of the authentic-donor (Figure 1Ci), with deletions ranging from 1 to 57 nts in length (Figure 1Cii). 89% of *AM*-variants result in use of a single cryptic-donor, 9% activate 2 cryptic-donors and 2% activate 3 or more cryptic-donors (Figure 1Ciii).

**B)** *‘Cryptic-Modified’ (CM):* a genetic variant modifies a cryptic-donor and does <u>not</u> affect the authentic-donor (n = 2,252) (Figure 1Bii). *CM*-variants most frequently affect the D^+2^ position of the cryptic-donor (Figure 1Di), with 32% of all *CM*-variant single nucleotide variants (SNVs) changing the cryptic-donor essential splice motif from GC to GT (Figure 1Dii).

***C)*** *‘Authentic-Modified / Cryptic-Modified’ (AM/CM)*: a genetic variant that simultaneously modifies the authentic-donor <u>and</u> nearby cryptic-donor (n = 373) (Figure 1Biii). *AM/CM*-variants also most frequently affect the D^+2^ position (122/373) of the cryptic-donor (Figure 1Ei), with 31% of SNVs converting a GC to GT (Figure 1Eii). Deletions range from 1 to 36 nts in length (Figure 1Eiii)

### 87% of activated cryptic-donors lie within 250 nt of the authentic-donor, with an average of 36 other decoy-donors (any GT or GC) within this distance not used by the spliceosome

95% of cryptic-donors activated by *AM*-variants and 65% of cryptic-donors activated by *CM*-variants, lie within 150 nt of the authentic-donor (Figure 2A, B). By definition, *AM/CM*-variants activate a cryptic-donor that spatially overlaps the authentic-donor; 26% of *AM/CM* cryptic-donors lie at either the E^−4^ or D^+5^ position (Figure 2C, bottom). For exonic cryptic-donors activated at E^−4^, the GT at D^+1/+2^ of the authentic-donor becomes D^+5/+6^ of the cryptic-donor; conversely for intronic cryptic-donors activated at D^+5^, the GT at D^+5/+6^ of the authentic-donor becomes D^+1/+2^ of the cryptic-donor) (Figure 2C, top).

**Fig 2.**
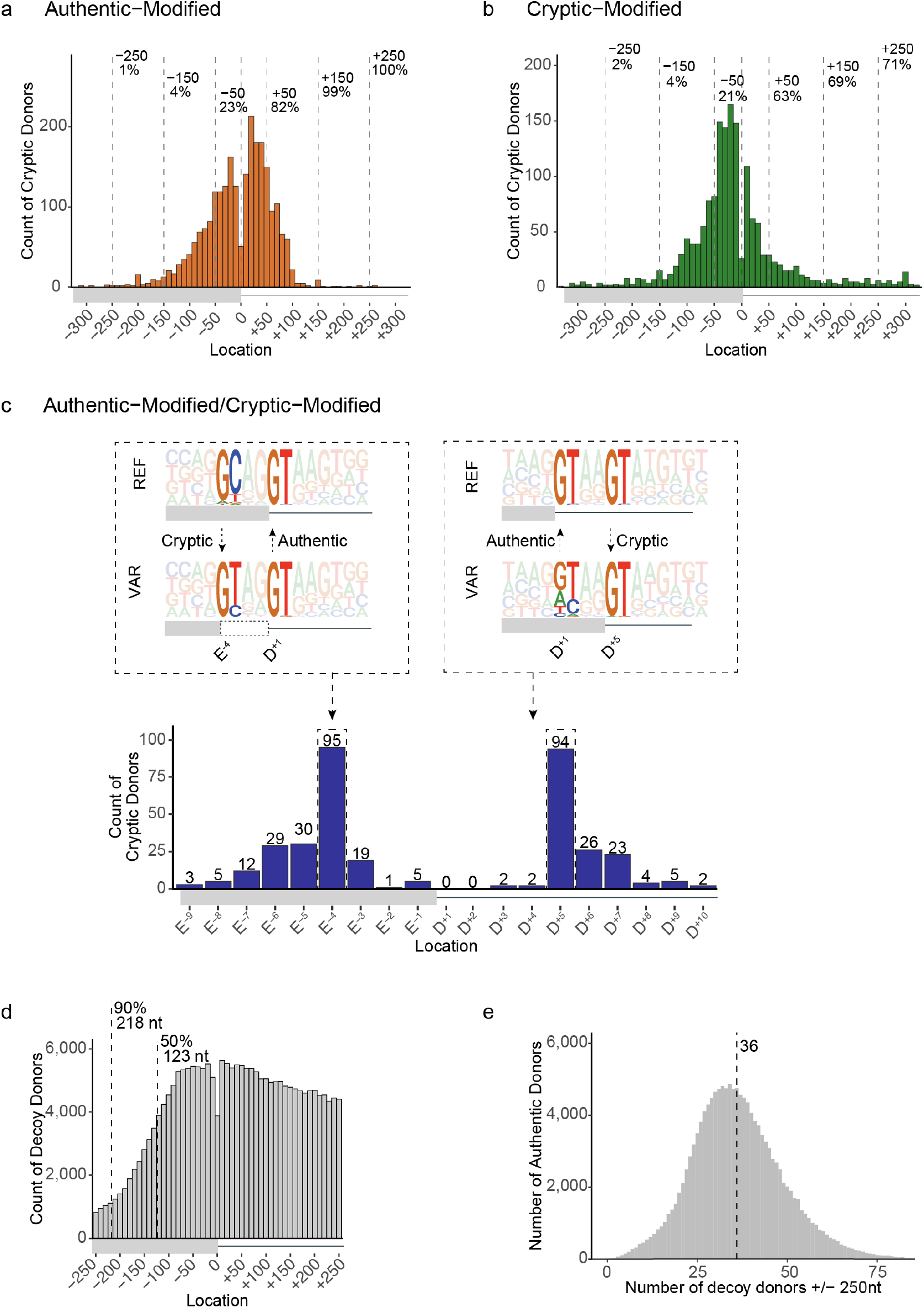
Cryptic-donor activation is influenced by proximity to the authentic-donor. **a-b)** Distribution of cryptic-donors activated by **(a)** *AM*-variants and **(b)** *CM*-variants. Location (x-axis) denotes the distance of the cryptic-donor from the authentic-donor, with negative values upstream into the exon and positive values downstream into the intron. **c)** (Bottom) Distribution of cryptic-donors activated by *AM/CM*-variants. (Top) Pictograms showing the Reference (REF) and Variant (VAR) sequences for *AM/CM*-variants. Activated cryptic-donors are prevalent at E^−4^ (left) and D+5 (right) due to conserved GTs at D^+1/+2^ and D^+5/+6^ of the conserved donor splice-site sequence. **d)** Frequency of naturally occurring decoy-donors (any GT or GC) +/− 250 nt of authentic-donors in the human genome (hg19). Dashed lines indicate the 50^th^ and 90^th^ percentile for exon length. The decline in exonic donors is due to relatively fewer longer exons. **e)** Distribution of the number of decoy-donors in the +/−250 nt surrounding each authentic-donor in the human genome. Dashed line shows that there are an average of 36 decoy-donors within 250 nt of each authentic-donor.

While decoy-donors are present everywhere, those selected as cryptic-donors by the spliceosome in the context of a genetic variant appears strongly influenced by their proximity to the authentic-donor (Figure 2A-B), as shown by their enrichment at proximal locations relative to all decoys present in the genome (Figure 2D). The steep decline in exonic decoys (Figure 2D, left) is due to the shorter lengths of exons limiting their frequency at these distances (50^th^ and 90^th^ percentile for exon length shown). Notably, each authentic-donor in hg19 has on average 36 decoy-donors within +/−250 nt <u>not</u> used by the spliceosome – indicating that features other than proximity to the authentic-donor define a usable cryptic-donor.

### Only 31-67% of cryptic-donors are scored as being stronger than the authentic-donor, with discordance between algorithms

We examined the ability of four algorithmic measures of ‘splice-site strength’ to predict cryptic-donor activation (Figure 3). We compared the performance of MaxEntScan (MES)^13^, NNSplice (NNS)^12^ and SpliceAI (SAI)^11^ as well as our own method termed ‘Donor Frequency’ (DF) (see methods and Figure S1 for details, Figure S2A-C for full plots). DF calculates the median frequency of four consecutive ‘windows’ of nine nucleotides in length (from E^−4^ to D^+8^) among all donors in hg19, converted to a cumulative frequency distribution. For example, if a E^−3^ to D^+6^ sequence has a raw frequency of 222, this combination of nine bases occurs at the analogous position for 222 annotated authentic-donors in hg19, corresponding to the 35^th^ percentile of a cumulative frequency distribution across the human genome (see Figure S1C). DF provides a barometer related to how common a given donor sequence is in humans, as a measure of splicing competence. For these and all further analyses, we excluded the 1,113 cryptic variants in the database derived from SAI predictions already validated on GTEx RNA-seq data^11^. Our nomenclature of REF and VAR corresponds to the reference (REF) or variant (VAR) donor sequence.

**Fig 3.**
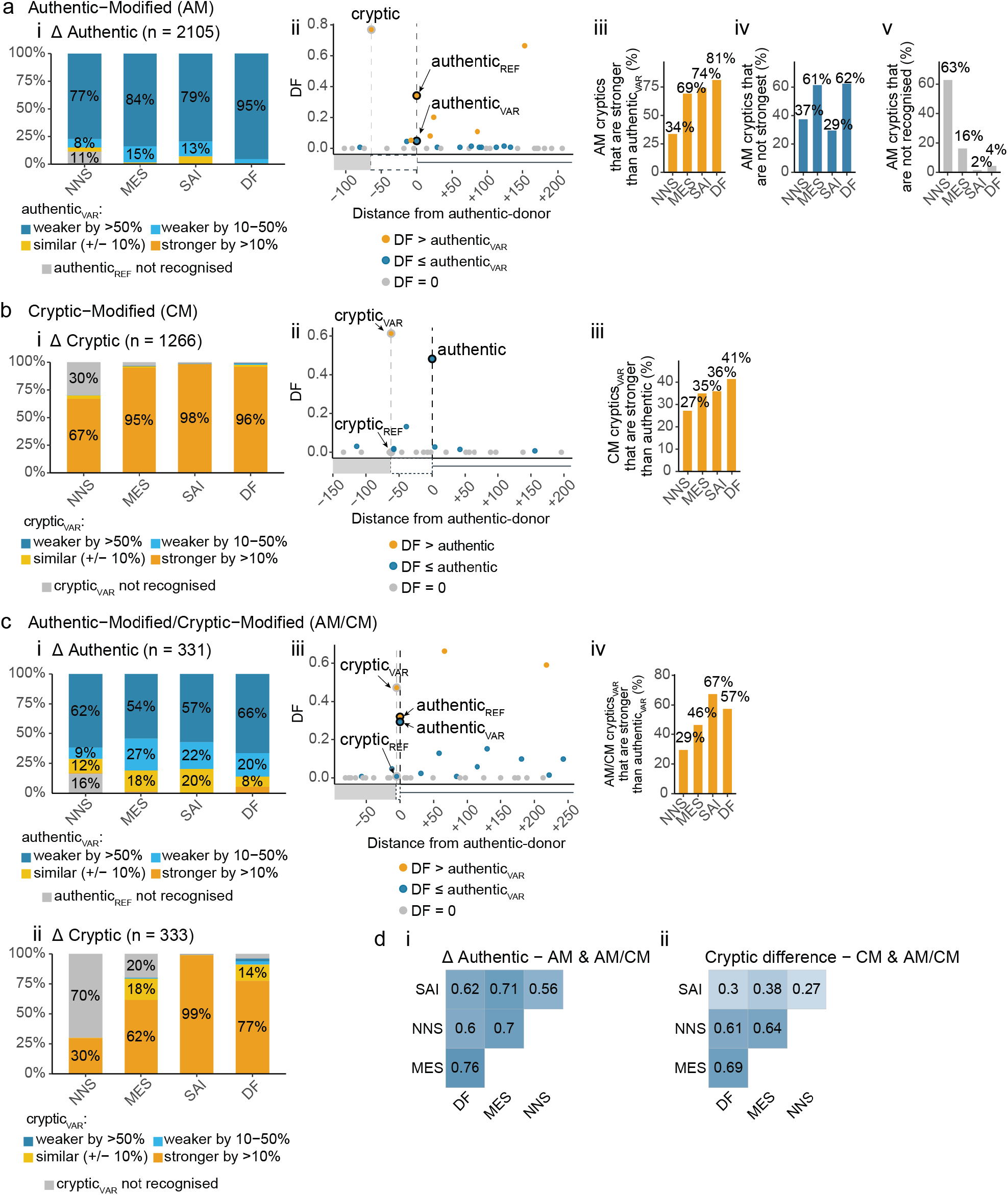
Cryptic donor activation is influenced by relative strength. **a)** Assessment of algorithmic scores of splice-site strength for *AM*-variants. ***i)*** Proportion of variants with authentic-donor Δ scores (Authentic_VAR_/Authentic_REF_) in each of the categories shown in the figure key. Most *AM*-variants weaken the authentic-donor by > 50% (*dark blue*). See Figure S2 for full plots. ***ii)*** Example variant showing the Donor Frequency (DF) scores (see Figure S1) for the cryptic-donor (DF = 0.77), versus the reference (REF = 0.34) and variant (VAR = 0.05) authentic-donor, as well as surrounding decoys not used. Vertical dotted lines indicate position of authentic- and cryptic-donors. Donors coloured according to the figure key. ***iii)*** Percent of cryptic-donors stronger than the authentic VAR. ***iv)*** Percent of cryptic-donors that are <u>not</u> the strongest donor splice-site within 250 nt. ***v)*** Percent of *AM*-variant activated cryptic-donors that are <u>not</u> recognised by each algorithm (i.e. score of 0). **b)** Strength measures for *CM*-variants. ***i)*** Proportion of variants with cryptic-donor Δ scores (Cryptic_VAR_/Cryptic_REF_) in each of the categories shown in legend. Most *CM*-variants strengthen the cryptic-donor by > 10% (*dark yellow*). See Figure S2 for full plots. ***ii)*** As in **a*ii*. *iii)*** As in **a*iii*. c)** Strength measures for *AM/CM*-variants. ***i)*** As in **a*i***. ***ii)*** As in **b*i*. *iii)*** As in **a*ii*. *iv)*** As in **a*iii*. d)** Pearson correlation of strength measures. ***i)*** Δ Authentic (VAR/REF) for *AM* & *AM/CM*-variants (all variants which affect the authentic-donor). ***ii)*** Cryptic difference (VAR – REF) for *CM* & *AM/CM*-variants (all variants which affect the cryptic-donor).

For *AM*-variants, activation of a cryptic-donor typically occurs in the context of a variant that weakens the authentic-donor by > 50% (Figure 3Ai, *dark blue*). While many *AM* cryptics are stronger than the authenticVAR (Figure 3Aiii, example shown in Figure 3Aii), a substantial subset are <u>not</u> the strongest decoy-donor within 250nt (Figure 3Aiv). In fact, many activated cryptic-donors are not recognised as *bona fide* donor by the respective algorithms, notably NNS (Figure 3Av).

Intuitively, for most *CM*-variants the cryptic is strengthened by 10% or more by the variant (Figure 3Bi, *orange*, example shown in Figure 3Bii). Counter-intuitively, less than half of activated cryptics_VAR_ outcompete the authentic (Figure 3Biii). Along similar lines, intuitively, for a majority of *AM/CM*-variants the authentic is weakened (Figure 3Ci, *blue)* while the adjacent cryptic is strengthened by the variant (Figure 3Cii, *orange*, example shown in Figure 3Ciii). Counter-intuitively, only 29 - 67% of AM/CM-cryptics_VAR_ outcompete the authentic-donor_VAR_ (Figure 3Civ). Despite similar overall performance for each algorithm, they showed discordance in variant outcome predictions (Figure 3D) and measures of splice-site strength (Figure S2D).

In summary, four independent algorithms concur that cryptic-donor activation typically occurs in response to weakening of the authentic-donor (85 - 99 % weaker) or strengthening of the cryptic-donor (67 - 98% stronger). However, only 35 - 70% of activated cryptic-donors are stronger than the authentic-donor_VAR_, and for unmodified cryptic-donors, 29 - 62% are not the strongest decoy-donor within 250 nt. Thus, while relative strength of the authentic- and cryptic-donor influence spliceosomal use, there are other factors at play.

### Competitive decoy-donors are specifically depleted proximal to the exon-intron junction

Decoy-donors of comparable or greater strength to the authentic-donor rarely occur within 150 nt (Figure 4A, top, *red*). Exonic and intronic regions around donors have characteristic single and dinucleotide frequencies (Figure S3). In particular, the first 50 nt of the intron often shows enrichment in G and T dinucleotides, with distinct patterns: 1) G repeats are enriched in the shortest of introns and T repeats in the longest (Figure S3C); 2) Introns with G (or C) at the D^+3^ position are enriched in G dinucleotides whereas introns with A (or T) at the D^+3^ position are enriched in T dinucleotides (Figure S3D); 3) Introns with rare donors (low DF) are enriched in T-repeats compared to introns with the most common donors (Figure S3E). Therefore, we adapt a method^18^ to assess if decoy-donors occur more or less commonly than expected by random chance, given these and any other dinucleotide preferences (see Methods and Figure S4).

**Fig 4.**
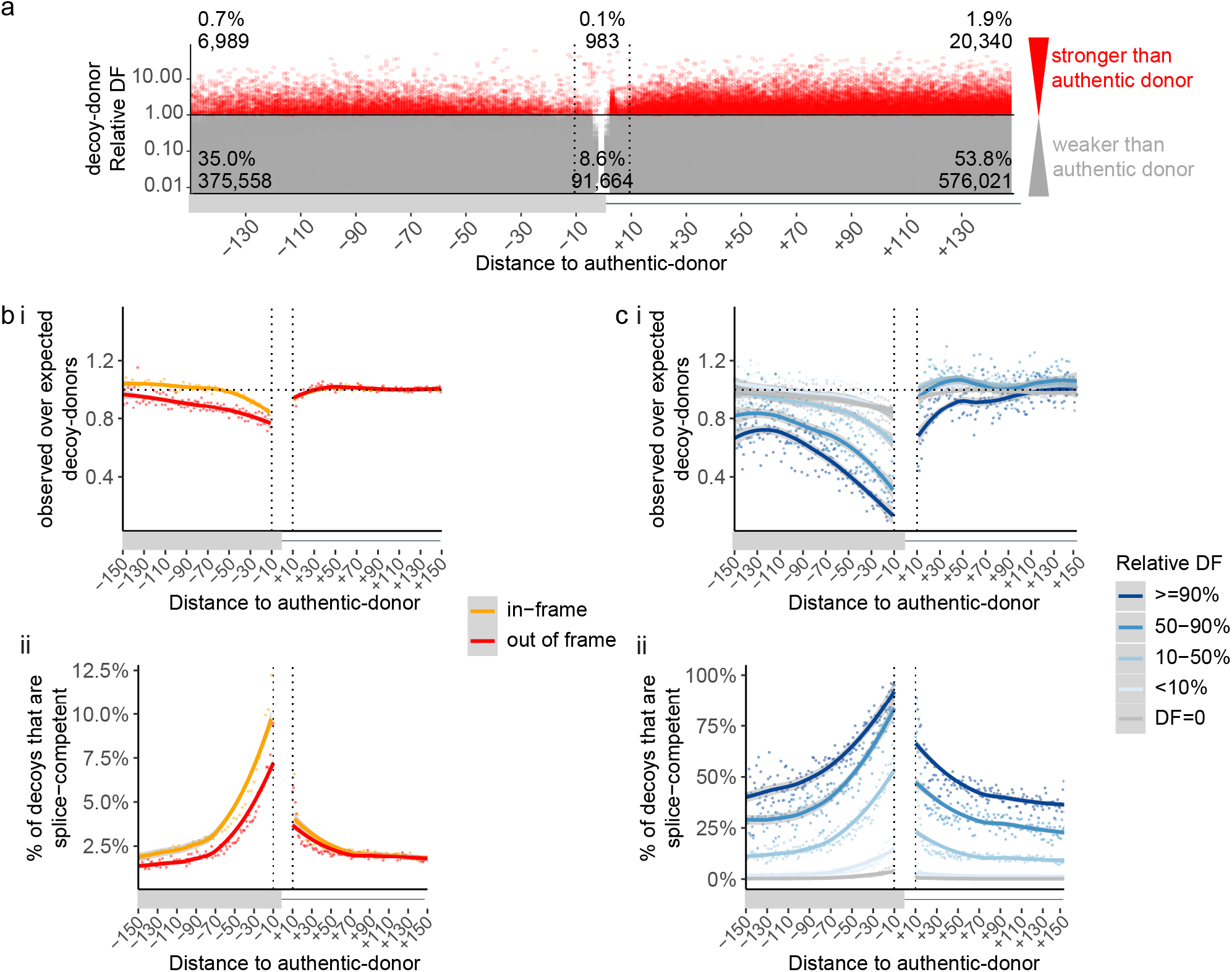
Competitive decoy-donors are specifically depleted proximal to the exon-intron junction. **a)** The relative DF of all decoy-donors within 150 nt of authentic-donors (decoy-donor DF / authentic-donor DF). Decoy-donors with a stronger DF score than the authentic-donor are shown in *red*, otherwise *grey*. **b*i)*** Depletion of ‘GT’ decoy-splice sites (observed/expected) (see Methods and figure S4). Exonic donors where use of the decoy-donor would be in-frame are shown in *orange*, whereas those out-of-frame (or intronic) are shown in *red*. ‘GT’ decoy-donors show increasing exonic depletion approaching the authentic-donor, and negligible depletion in the intron. **b*ii***) S*plice competence* of in-frame and out of frame ‘GT’ decoy-donors, defined as rare spliceosomal use in aggregate splice-junction data from 40,233 publicly available RNA-seq samples. At each distance from the authentic splice-site, the number of decoy-donors with evidence of *splice competence* (i.e. at least 1 splice-junction read in aggregate RNA-Seq data) is divided by the total number of naturally occurring decoy-donors at that position. A higher proportion of decoy-donors are *splice competent* at locations more proximal to the authentic-donor **c*i)*** depletion of ‘GT’ decoy-donors as in **b*i***, split according to decoy-donor DF relative to the authentic-donor (decoy-donor DF / authentic-donor DF). There is negligible depletion of decoy-donor sequences that do not exist as a bonafide donor in hg19 (DF = 0, *grey*), with increasing depletion of exonic decoy-donors closer in DF to the authentic donor (*blue gradient*). ***ii)*** S*plice competence* of ‘GT’ decoy-donors as in **b*ii***, split as in **c*i***. *Splice competence* of decoy-donors is influenced by their relative DF <u>and</u> proximity to the authentic splice-site.

‘GT’ decoy-donors show increasing exonic depletion approaching the authentic-donor, with out-of-frame decoys (*red*) depleted more than in-frame decoy-donors (*orange*), while showing negligible depletion in the intron (Figure 4Bi). ‘GC’ donors show no depletion in either the exon or the intron (Figure S5A).

We next assessed what proportion of decoy-donors in the genome are used, albeit rarely, via unannotated splice-junctions detected across 40,233 publicly available RNA-seq samples (see methods). Unannotated splice-junctions (representing stochastic mis-splicing), seen rarely in RNA-seq samples aggregated across a population, constitute functional evidence that both splicing reactions can be executed using a decoy-donor: hence we term these donors *’splice competent*’. The remaining decoy-donors, for which no RNA-Seq splice-junctions are detected, we term ‘*non-splice competent*’ – acknowledging this assertion is confounded by low read depth for many transcripts.

The proportion of exonic *splice competent* ‘GT’ decoy-donors (relative to all decoys) dramatically increases with proximity to the authentic-donor, with intronic decoys showing only a modest change (Figure 4Bii), mirroring depletion patterns (4Bi). Decoy-donors closer to the authentic-donor appear to be inherently more *splice competent*, supported by the fact that they are depleted in the human genome. Less than 4% of exonic ‘GC’ decoy-donors show evidence of *splice competence*, even very close to the exon/intron junction, in line with the observed lack of depletion (Figure S5B).

The ability of DF to measure strength/competition of a splice-site sequence is evidenced by Figure 4C. While there is negligible depletion of decoy-donor sequences that do not exist as a *bona fide* donor at any exon-intron junction in hg19 (DF = 0, *grey*), there is clear depletion of exonic decoy-donors closer in DF (50-90% DF; Figure 4Ci, *mid-blue*), or of similar or greater DF (≥ 90% DF, *dark blue*) (Figure 4Ci, left), relative to the authentic-donor. Interestingly, except for the most competitive decoy-donors (≥ 90% DF; Figure 4Ci, right, *dark blue*), decoy-donors show no depletion in the intron. Concordantly, we see increasing *splice competence* of decoy-donors at the end of the exon with increasing relative DF, and to a lesser extent at the start of the intron (Figure 4Cii).

### Why do intronic decoy-donors show lower splice competence?

The lack of depletion and lower *splice competence* of decoy-donors in the intron (Figure 4B-C) was initially perplexing, given that cryptic-donors are just as common in the intron as in the exon. However, we reasoned distinctive nucleotide preferences in the first 50 nt of the intron could affect measures of depletion, and/or, influence the ‘usability’ of decoy-donors in this region. For example, G-repeat splicing regulatory elements (SREs) are abundant within the first 50 nt of the intron^19–21^.

Defining ‘competitive’ decoy-donors as those with a DF of at least 10% that of the associated authentic-donor (see Figure 4C), in the first 50 nt of the intron, competitive decoy-donors overlapping G-triplets show no depletion and indeed appear enriched (Figure 5Ai, intron- *orange*). In contrast competitive decoy-donors <u>not</u> overlapping G-triplets are depleted (Figure 5Ai, intron- *grey*). Additionally, intronic decoy-donors <u>not</u> overlapping G-triplets show greater splice competence that those overlapping G-triplets (Figure 5Aii, intron- *grey*). The reciprocity in these data is consistent with a masking effect of intronic G-repeat motifs on (strong) decoy-donors, likely due to RNA secondary structure and/or RNA binding proteins preventing their use.

**Fig 5.**
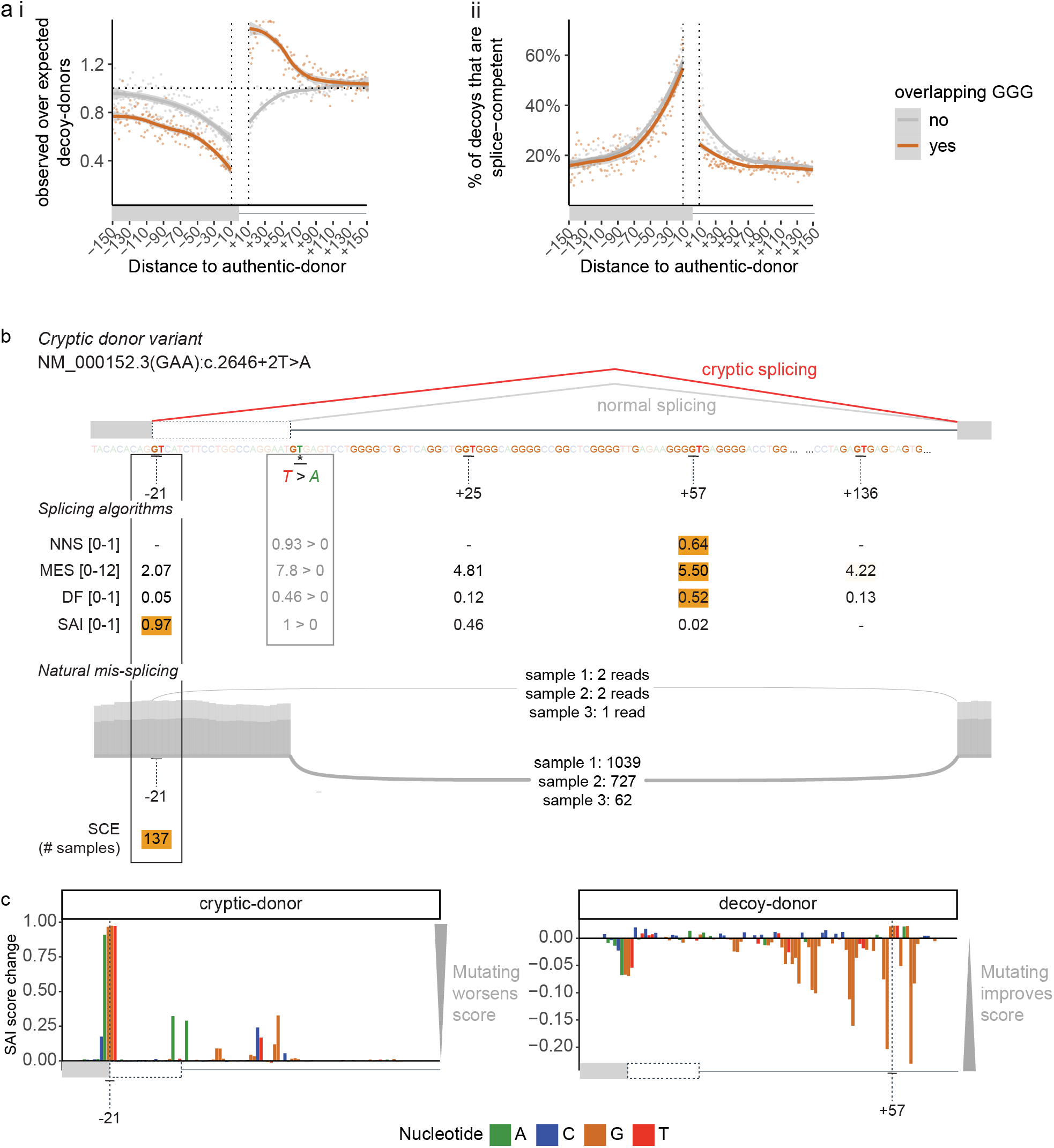
Utility of aggregate RNA-Seq splice-junction data to identify *splice competent* decoy-donors. **a)** Depletion *(**i**)* and *splice competence (**ii**)* of ‘GT’ decoy-donors that do or do not overlap G-triplets. Decoy-donors overlapping G-triplets are depleted in the exon but not in the intron, where they show enrichment because: *1)* G repeats are enriched in the first 50 nt of the intron (see Figure S3b) and *2)* donor sequences are commonly G-rich. Plots are limited to decoys with relative DF > 0.1 (defined as ‘competitive’ with the authentic-donor, see Figure 4C). **b)** Top: Schematic of an *AM*-variant identified in an individual with glycogen storage disease type II associated with *GAA* (NM_000152.3:C.2646+2T>A)^22^ with algorithm scores for authentic- (REF > VAR) decoy- and cryptic-donors. The strongest donor for each algorithm (score range shown in square brackets) is coloured *orange*. Below: Sashimi plot from three GTEx RNA-seq samples identifying use of the −21 cryptic-donor as a ‘*splice competent event’* (SCE, at least 1 read is detected in 137/30753 samples with detectable expression of *GAA*). SpliceAI (SAI) correctly scores the −21 cryptic-donor as the most likely cryptic-donor. **c)** Result of SAI *in silico mutagenesis* showing the bases contributing to predicted strength of the −21 cryptic-donor (left) and +57 decoy-donor (right). ‘SAI score change’ denotes the decrease (if positive) or increase (if negative) on the predicted strength of the donor when that nucleotide is mutated (see methods). Note that the presence of the cryptic-donor at −21 and intronic G-repeats negatively impact the score of the +57 decoy-donor according to SAI.

Figure 5B shows an example variant in *GAA* (NM_000152.3:C.2646+2T>A) identified in an individual affected with glycogen storage disease type II^22^ that induces cryptic splicing to a exonic donor 21 nt upstream of the authentic-donor. NNS, MES, and DF rank the decoy-donor at +57 as the most competitive donor - however this donor is enveloped within a G-repeat rich region which may mask it. Mis-splicing data finds no evidence of rare spliceosomal use of the +57 decoy. SAI does not predict the decoy at +57 to be a functional splice-site, instead predicting the cryptic-donor at −21 to be a functional splice site. Notably, natural mis-splicing data shows rare, natural spliceosomal use of the cryptic-donor at −21 in 137 samples, providing evidence that despite its weak primary motif, it is *splice competent* (Figure 5B).

SAI *in silico* mutation of the cryptic-donor at −21 and decoy-donor at +57 show that SAI deep-learning perceives the negative impact of the G-repeats on the usability of the +57 decoy-donor (Figure 5C). Intronic G-repeats are known examples of SREs that can influence splicing decisions^19,23^ (see Figure 5 and additional examples Figure S5C,D). The impact of extended sequence context on splicing decisions likely explains the inadequacy of current measures of splice site strength to effectively rank likely cryptic-donor(s) among all decoy-donors (any GT or GC) not positioned correctly at the exon-intron junction.

### 90% of confirmed cryptic-donors in *AM*-variants are detected as rare splice-junctions in RNA-Seq data, with 91% of unused decoy-donors absent

We assessed whether evidence of *splice competence* from RNA-seq data provides a viable means to prioritise cryptic-donors likely to be activated in the context of a genetic variant affecting the authentic-donor (i.e. *AM*-variants). Strikingly, 90% of *AM*-variant activated cryptic-donors are detected as *splice competent* events in aggregate RNA-Seq splice-junction data, while 91% of unused decoy-donors are absent. We took the top 4 *splice competent events* at each splice-junction (or all events if there were less than 4 detected) as our predicted cryptic-donors and this yielded a sensitivity of 87% and specificity of 95% (Figure 6Ai). Our method had a higher sensitivity than all four algorithms assessed (Figure 6aii, Figure S6A).

**Fig 6.**
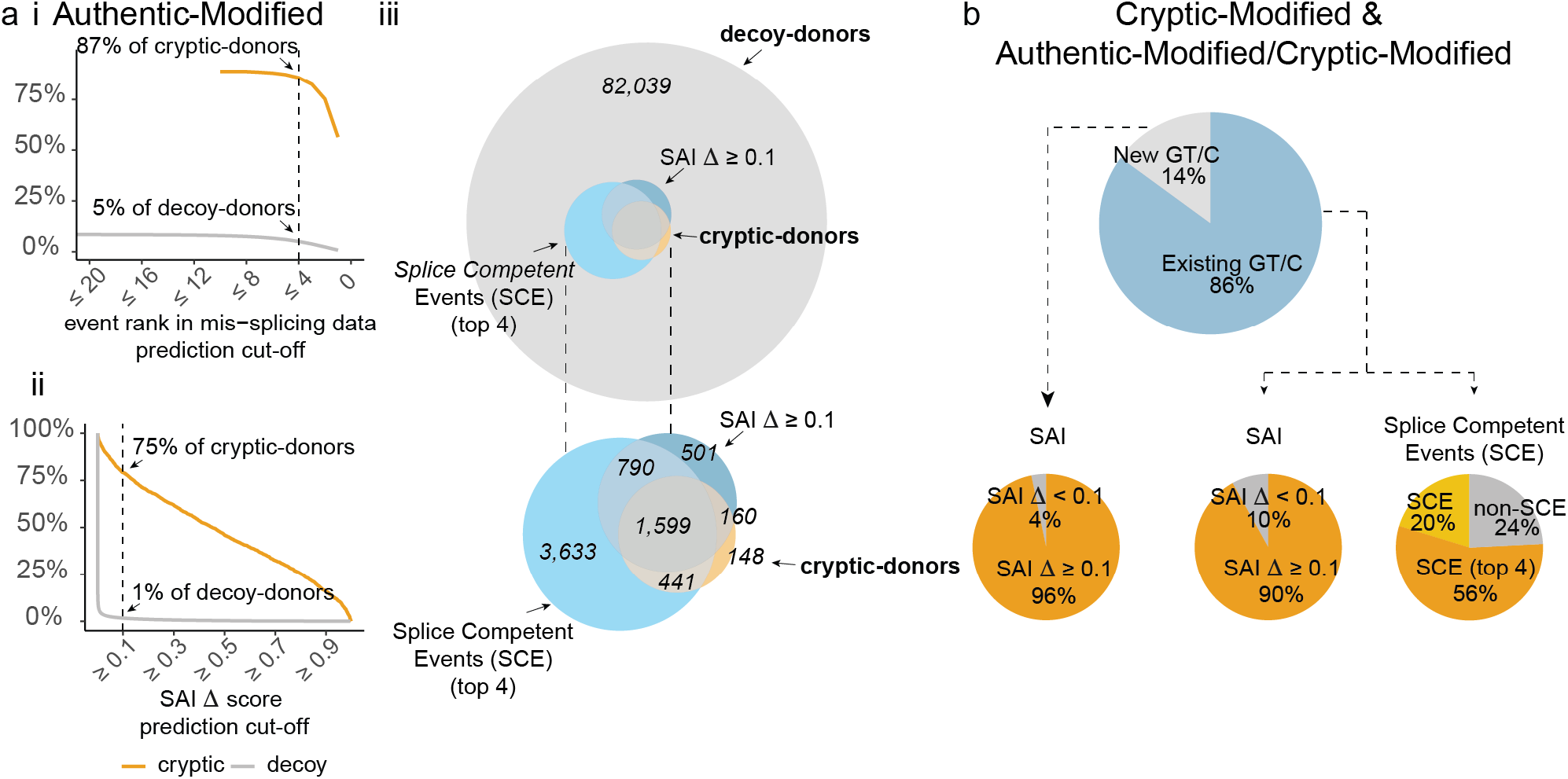
Ranking unannotated splice-junctions detected in aggregate RNA-Seq data potently informs cryptic-donor activation. *a)* The percentage of cryptic-donors correctly predicted, and decoy-donors incorrectly predicted at different cut-offs for *(i)* event rank in aggregate mis-splicing data and *(ii)* SAI Δ score (donor_VAR_ / donor_REF_). **i**) The dotted line denotes cryptic- and decoy-donors predicted using a cut-off of events ranked 4 or less. 87% of cryptic-donors activated by *AM*-variants are among the top 4 unannotated splice-junctions within 250 nt of the authentic-donor, with 95% unused decoy-donors absent in splice-junction data. *ii)* While SpliceAI (SAI) outperforms other algorithms using a cut-off of Δ scores 0.1 and above (see figure S6A), it predicts only 75% of cryptic-donors (SAI Δ ≥ 0.1) and accurately excludes 99% decoy-donors unused. ***iii)*** (top) Venn diagram showing the overlap of cryptic-donors with those predicted by SAI or our SCE (*splice competent events*) method, among the entire pool of decoy-donors. (bottom) magnification of internal Venn. **b)** Cryptic-donors activated by *CM*- and *AM/CM*-variants. (top) ‘New GT/C’ (Light *grey*) denotes variants creating a GT or GC essential splice-site motif (SCE is inherently unsuitable for these variants). ‘Existing GT/C’ (*blue*) denotes variants that modify a decoy-donor with a pre-existing GC or GT essential splice-site. (bottom) For *splice competent* events, the *orange* segment denotes cryptic-donors in the top 4 *splice competent* events, *yellow* segment denotes where the cryptic-donor is present as a *splice competent* events but is not in the top 4 events, and *grey* denotes cryptic-donors absent in aggregate RNA-Seq splice-junction data (non-SCE i.e. no prior evidence of being a *splice competent event*).

Therefore, splice-junction data provides potent predictive information with respect to both true positives (cryptic-donors) and true negatives (decoy-donors). Most cryptic-donors activated are observed as very low frequency splice-junctions (44% had a maximum of 4 reads or less, Figure S6B). Sensitivity of using RNA-Seq splice-junction data is inherently influenced by read-depth of the target transcript: more than 85% of cryptic-donors are detected in genes with a read depth of > 250 for the normal exon-exon splice-junction; whereas only 29% of cryptic-donors are detected in mis-splicing data for genes where normal splicing had a maximum read count of < 100 (Figure S6C).

We define SAI prediction of cryptic-donor activation as a donor Δ score of 0.1, which accurately predicts 75% of cryptic-donors and inaccurately predicts only 1% of decoy-donors (Figure 6Aii). SAI shows the lowest predictive accuracy for more distal cryptics - only 55% of cryptic-donors further than 100 nt from the authentic splice site had a Δ score above 0.1 (Figure S6D). When used in combination, SAI and RNA-Seq *splice competent* evidence accurately predict 2210/2389 (93%) of cryptic-donors (Figure 6Aiii) and inaccurately predict 6% of unused decoy-donors.

Use of RNA-Seq *splice competent* events has caveats for *CM*-variants and *AM/CM*-variants, particularly for variants that create a GT (or GC) motif, as an existing essential splice-site is requisite for stochastic mis-splicing. However, for the subset of *CM*-variants and *AM/CM*-variants where the variant modifies the extended splice site region of an extant GT/C decoy-donor (1525 variants, Figure 6B, top-*blue*), 76% are detected as rare splice-junctions in RNA-seq data, with 56% in the top 4 splice competent events.

D^+2^ GC>GT variants commonly activate cryptic-donors (Figure 1). However, because ‘GC’ donors are inherently much less *splice competent* than ‘GT’ donors (Figure S5B) they are infrequently seen in mis-splicing data, limiting predictive ability for these variants. For D^+2^ CM-variants, only 32% of the cryptic-donors are present in the top 4 mis-splicing events, as compared to 85% of cryptic-donors detected in the top 4 events for E^−3^ variants (Figure S6E; E^−3^ is the third to last exonic base). In short, even if a ‘GC’ decoy-donor shows no evidence of *splice competence*, conversion by a variant to a ‘GT’ donor presents high risk for cryptic-activation. SAI performed well for *CM*-variants and *AM/CM*-variants, correctly predicting 96% of variants which created an essential donor motif and 90% which modified an existing essential motif (Figure 6B).

## Discussion

The ultimate goal of splicing predictions is to determine if and how a genetic variant will induce mis-splicing of pre-mRNA. Consideration of the likely consequences of an essential splice-site variant is critical for pathology application of the ACMG-AMP code PVS1^15^ (null variant due to presumed mis-splicing of the pre-mRNA) and of equal importance to strategise functional testing for RNA diagnostics^14^. While activation of a cryptic-donor 6 nucleotides away will remove or insert two amino-acids, activation of a cryptic-donor 4 nucleotides away will induce a frameshift, with vastly different implications for pathology interpretation.

We learned five key lessons from our analyses of 4,811 variants in 3,399 genes: **1)** Decoy-donors shown to be *splice competent* via aggregate RNA-Seq splice-junction data have the greatest probability of activation as cryptic-donors. **2)** Proximity to the authentic splice site increases likelihood of spliceosomal mis-use (*splice competence*) of a decoy-donor. **3)** Cryptic-donors do not necessarily need to be ‘stronger’ than the authentic-donor to present substantive risk for mis-splicing. **4)** Intronic G-repeats diminish the likelihood of spliceosomal recognition and use of intronic decoy splice sites. **5)** SAI’s deep-learning appreciates the broader sequence context determining spliceosomal ‘usability’ of a decoy splice-site, including 2-4 above, though less accurately predicts activation of more distal decoy-donors (>100 nt from the authentic-donor).

SAI’s deep learning presents a major improvement in predicting cryptic-donor activation. However, use of SAI in a pathology context is limited by the challenge of deriving a clinically-meaningful interpretation of a number on a 0 – 1 scale, without independently verifiable evidence. In contrast, aggregate RNA-Seq splice-junction data provides an accurate, evidence-based means to rank cryptic-donors likely to be activated by genetic variants.

Brandão et al.^24^ used deep sequencing of twelve major cancer susceptibility genes to catalogue all alternative and aberrantly spliced transcripts. They found variant-activated cryptic splicing was often seen at much lower levels in disease controls, suggesting that the dominant transcript in rare disease may be seen as a stochastic mis-splicing event in other samples. Herein we use the breadth of publicly available RNA-seq data across numerous tissues to capture stochastic mis-splicing events across all genes, to comprehensively catalogue all *splice competent* cryptic splicing events. Prospectively, the potency of RNA-Seq *splice competent* data can be enhanced by ultra-deep sequencing.

When used in combination, SAI and RNA-Seq *splice competent* evidence accurately identify 93% of cryptic-donors (true positives) and inaccurately identify 6% of unused decoy-donors (false positives). The heightened sensitivity of our method is of vital importance for pathology assessment of variants affecting the essential donor splice-site: as not considering a likely cryptic-donor activated can lead to profound complications in variant interpretation. It is also easy to envisage extending the method to predict other mis-splicing events such as exon skipping, and an analogous formulation for the prediction of mis-splicing events at the acceptor splice site.

RNA-Seq *splice competent* data can reliably identify distal cryptic-donors with high likelihood of activation, which may not be identified by SAI. Conversely, SAI can reliably identify cryptic-donors with high likelihood of activation not detected in RNA-Seq *splice competent* data, due to low read depth of the target gene.

Importantly, RNA-Seq empirical evidence of *splice competence* can augment pathology consideration of probable consequences of splice site variants on pre-mRNA splicing (and the encoded protein), to assist clinical decision-making and the diagnostic classification of variants.

## Methods

### Creating a database of cryptic-donor variants

Variants were derived from several sources: 1) 439 variants curated from literature, predominantly comprised of 364 variants in DBASS5^16^ and supplemented by curation from published literature of 75 additional variants^25,26^ 2) 4,372 variants derived from RNA-seq studies: Variant-associated aberrant cryptic-donor activation detected from RNA-seq data identified by SAVnet in somatic tumor samples (n = 3,259)^17^ and 1,113 variants identified in GTEx samples by spliceAI and verified using RNA-seq data^11^. The following inclusion criteria applied: 1) Variants had to occur within E^−4^-D^+8^ of the authentic or the cryptic-donor, otherwise they were excluded as outside the bounds of this analysis. 2) annotated cryptic-donors were within the same exon/intron as the variant (i.e. between the 5’ end of the exon and 3’ end of the intron surrounding the affected donor). 3) The annotated cryptic-donor ALT sequence had to have an essential GT/GC dinucleotide at D^+1^/D^+2^, to minimise misannotated variants being included.

#### i. Annotating variant categories

We annotated variants with categories we defined– if the variant occurred within E^−4^-D^+8^ of the authentic-donor, it was an *AM*-variant, if it occurred within E^−4^-D^+8^ of the cryptic-donor it was a *CM*-variant, and if it occurred within E^−4^-D^+8^ of both the authentic- and cryptic-donor it was an *AM/CM*-variant. For 37/373 of AM/CM-variants, an additional unmodified cryptic-donor was activated, in addition to the cryptic-donor modified by the variant- these were excluded from analyses.

#### ii. Sequence extraction

We used the 1000genomes Phase2 Reference Genome Sequence (hs37d5) version of hg19^27^ implemented in RStudio to extract a window of (up to) 500nt of genomic sequence centred on the authentic-donor for each variant. For each variant in the cryptic-donor database, we extracted up to 250 exonic nucleotides in the 5’ direction (i.e. if the exon was only 50 nt the window of analysis would be 50 nucleotides), and up to 250 intronic nucleotides in the 3’ direction, in the same fashion (Figure 1A).

#### iii. Finding authentic, cryptic & decoys

From the (up to) 500 nt of sequence we pulled E^−4^-D^+8^ sequences for the authentic- and cryptic-donor before and after each variant (REF and VAR respectively). We also identified any other essential donor dinucleotides (i.e GT or GC) which were present in the sequence and extracted the E^−4^-D^+8^ sequence surrounding them. These sequences we define as ‘decoy’-donor- sequences containing the essential donor dinucleotides (i.e. a GT or a GC) but which *weren’t* utilised by the spliceosome as a result of the variant (Figure 1A, lower). For intronic decoy-donors, we excluded any which would result in an intron too short to be spliced (as defined by the 5^th^ percentile for intron length in the human genome)^28^. Importantly, without additional filtering, no cryptic-donors in the database violated this rule.

### Algorithms for splice site strength

We retrieved predicted scores for authentic-donors, cryptic-donors and decoy-donors in the database in both the REF and VAR sequence context, for four algorithms. **1)** MaxEntScan (MES)^13^ scores were retrieved using the perl script provided at http://hollywood.mit.edu/burgelab/maxent/Xmaxentscan_scoreseq.html. MES scores below 0 were standardised to 0 as predicted ‘non-functional’ splice sites **2)** NNSplice (NNS) ^12^ scores were retrieved using the online portal (https://www.fruitfly.org/seq_tools/splice.html), set to human, with default settings (i.e. a minimum score of 0.4, with any scores below predicting a ‘non-functional’ splice site) **3)** SpliceAI (SAI)^11^ scores were retrieved using a script adapted from the SAI github repository (https://github.com/Illumina/SpliceAI) which allows spliceAI to score custom sequences. We rounded the scores to three decimal places, and scores at 0 when rounded as such were termed ‘non-functional’ splice site predictions. **4)** Donor Frequency (DF) calculates the median frequency (in hg19) among four 9 nt windows of sequence spanning the donor (see Figure S1B-D) converted to a cumulative percentile distribution scale. DF provides a barometer related to how common a given donor sequence is in humans, as a measure of splicing competence. An example of a DF calculation is shown in figure S1D, where a median DF raw value of 179 lies at the 31^st^ percentile of a cumulative frequency distribution. After assessing several window lengths (figure S1B) we used 9nt windows as optimally encompassing the splice site (Figure S1C).

### Naturally occurring decoy-donors

Our set of naturally occurring human decoy-donors were derived from the set of all canonical Ensembl transcripts (Release 75)^29^, with first and last introns and single exon transcripts removed. We filtered the set so that junctions were within the open reading frame for the given gene, so we knew that cryptic splicing here would affect the protein. We also removed exons with alternative 5’ or 3’ ends already annotated in different transcripts. We used these criteria to form a set of 142,014 canonical exon-intron junctions that are not alternatively spliced (or at least not annotated as such). We extracted sequences surrounding authentic-donors and extracted all decoy-donors just as for the cryptic database (see methods section *creating a database of cryptic-donor variants)*

### Decoy-donor depletion

Decoy-donor depletion was calculated using an adapted method^18^ that controls for dinucleotide frequencies (Figure S5). For exonic sequences, we took up to 150 nt or the maximum length of the exon, whichever was shorter (and similarly for the intron, stopping 50nt from the 3’SS). We limited analysis to 150nt of exonic sequence as the majority of exons are shorter than this. We then shuffled exonic and intronic sequences separately, maintaining dinucleotide frequencies (using shuffle_sequences with euler method from universal motif R package^30^). We performed the shuffle 15 times and took the average count of decoy-donors at each nucleotide position as our ‘expected’ count at this position. The observed count of decoy-donors was then divided by the expected count at each position.

### Creating a database of splice competent events (SCE)

We had two sources of data for our database of *splice competent* events- RNA-seq data from Intropolis^31^ and GTEx ^32^. Intropolis is a set of ~42M exon-exon junctions found across 21,504 human RNA-seq samples from the Sequence Read Archive (SRA). Samples were aligned using Nellore et al.’s annotation-agnostic aligned Rail-RNA ^33^. Intropolis was downloaded from its dedicated github repository (https://github.com/nellore/intropolis). Per sample splice-junction files were obtained from GTEx (phs000424.v8.p2). Using Datamash^34^, splice-junction read counts were summarised across all samples for each unique splice-junction and translated from GRCh38 to GRCh37 using liftOver^35^.

For each set of junctions (Intropolis and GTEx), we cross-referenced and located junctions within Ensembl transcripts, keeping only events with at least one annotated splice site. We annotated cryptic-donor events by scanning for any unannotated donors used between the 5’ end of the exon and the 3’ end of the intron for that respective exon-intron junction, where the junction also spliced to the next annotated 3’SS. Then events from the two sources were merged and sample counts were tallied across the two datasets. If a splice-junction was seen in at least 3 samples, it was defined as ‘*splice competent’*.

### Sashimi plots

For figure 5B, and S6B-C sashimi plots were generated using 3 GTEx bam files for each example, each from the tissue with the highest TPM for the respective gene. Sashimi plots were created using ggsashimi ^36^.

### SpliceAI in silico mutagenesis plots

For figure 5C and S6B-C we performed *in-silico* mutagenesis as described in Jaganathan et al^11^. The importance score of each nucleotide was calculated as:

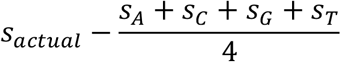

Where *s_actual_* is the score calculated on the genuine sequence, and *s_A_*, for example, is the score calculated when an A is substituted at this position.

## Acknowledgement

The Genotype-Tissue Expression (GTEx) Project was supported by the Common Fund of the Office of the Director of the National Institutes of Health, and by NCI, NHGRI, NHLBI, NIDA, NIMH, and NINDS. The data used for the analyses described in this manuscript were obtained from dbGaP accession number phs000424.v8.p2

## Author Contributions

Data curation and analysis: R.D. and H.J. Funding acquisition and supervision: S.T.C. Visualization: R.D. Writing – original draft: R.D. Writing review and editing: all authors.

## Competing Interests

S.T.C. and H.J. are named inventors of Intellectual Property (IP) described in part within this manuscript owned jointly by the University of Sydney and Sydney Children’s Hospitals Network. S.T.C. is director of Frontier Genomics Pty Ltd (Australia) who have licenced this IP. S.T.C. receives no payment or other financial incentives for services provided to Frontier Genomics Pty Ltd (Australia). Frontier Genomics Pty Ltd (Australia) has no existing financial relationships that will benefit from publication of these data. The remaining co-authors declare no conflicts of interest.

