## Supplemental Figures for "Empirical prediction of variant-associated cryptic-donors with 87% sensitivity and 95% specificity"

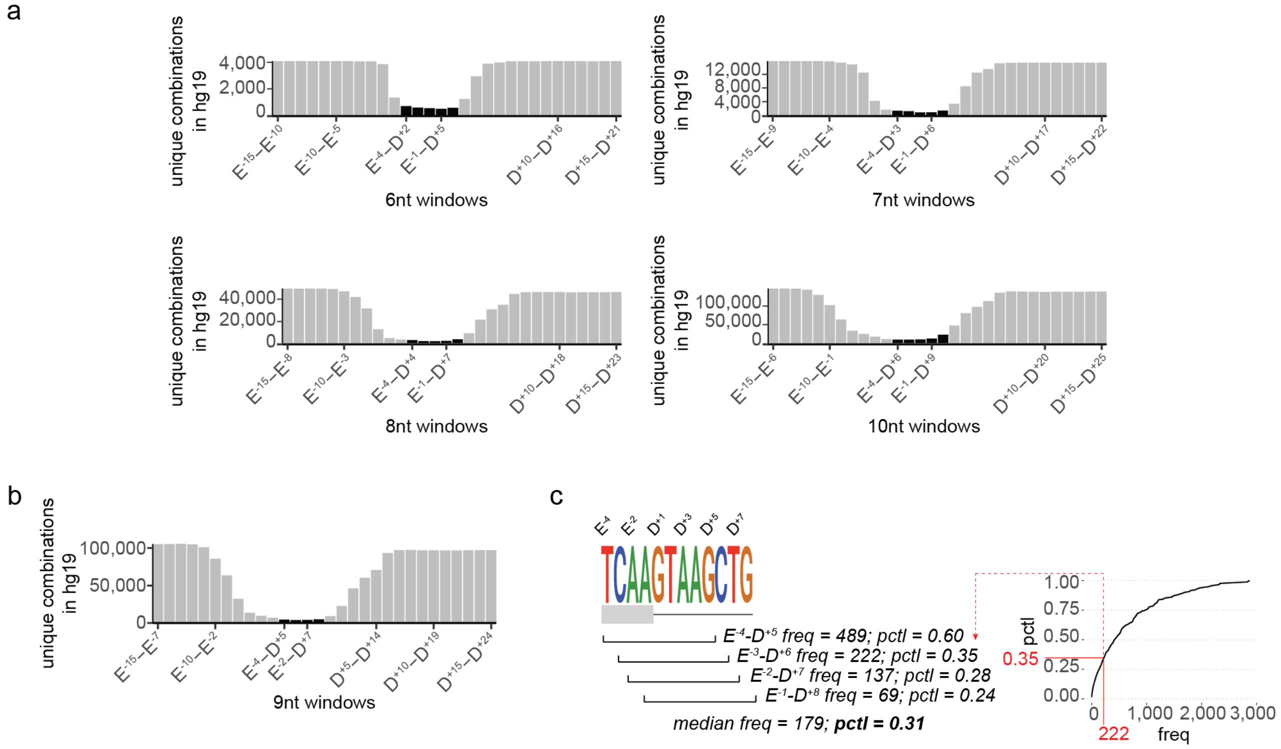


**Fig. S1 Calculation of Donor Frequency as a measure of donor strength. a-b)** Frequency of unique combinations of donor sequences at each position of the exon-intron junction in hg19, spanning 6, 7, 8, 10 **(a)** or 9 **(b)** consecutive nucleotides. Black bars denote windows overlapping the $E^{-4}$-$D^{+8}$ donor sequence window. Four sliding windows of 9 nt spanning the authentic-donor (coloured *black*), spanning 12 nt from the fourth-to-last exonic base ($E^{-4}$; E = exon) to the eighth intronic base ($D^{+8}$; D = donor),were used for DF calculation. **d)** Donor Frequency is calculated as the median frequency (in hg19) across each 9nt window, converted to a cumulative percentile distribution. DF provides a barometer related to 'how common' a given donor sequence is in humans, as a measure of splicing competence. In this example, a median DF raw value of 179 lies at the 31st percentile of a hg19 cumulative frequency distribution.

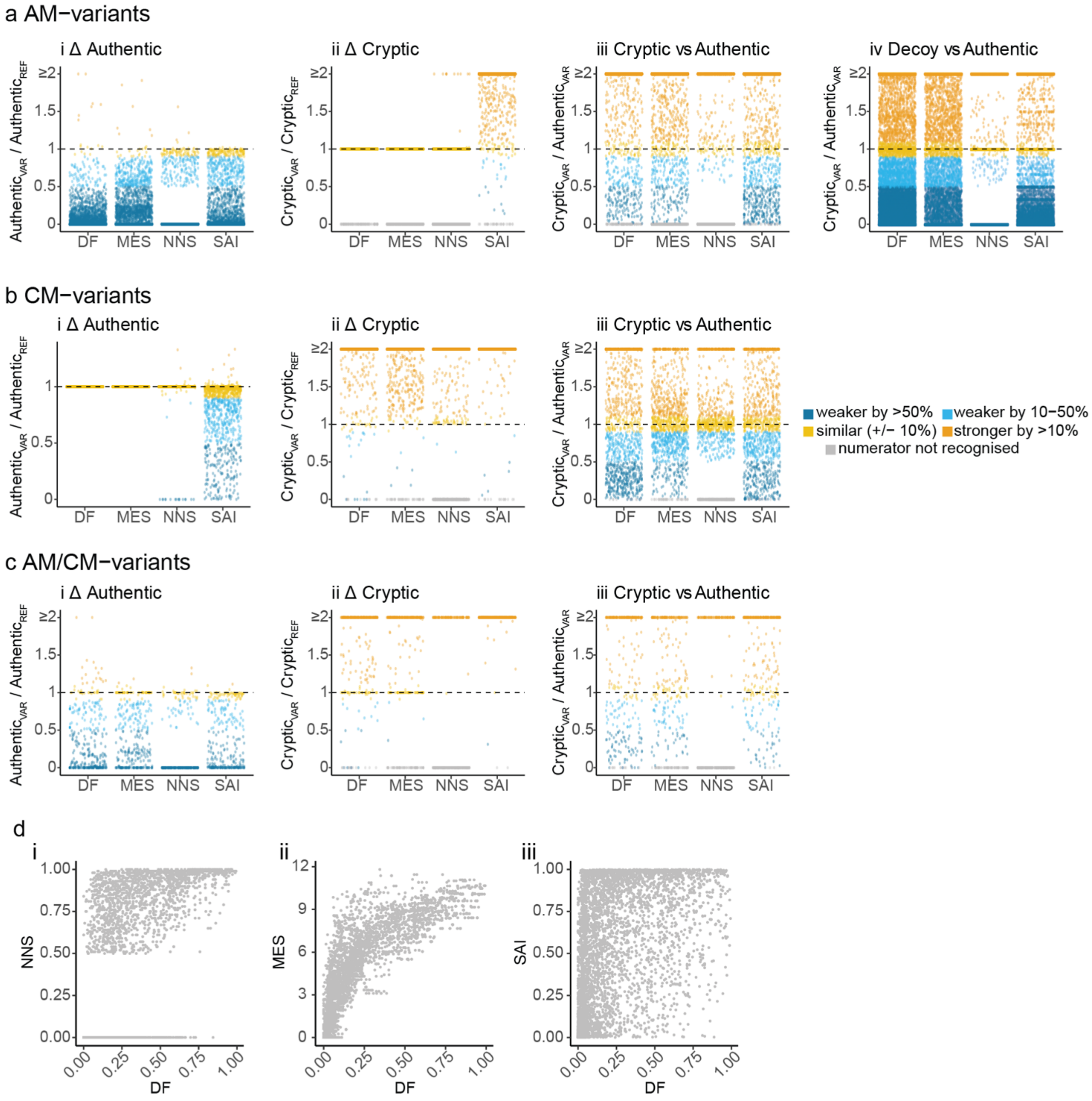


**Fig. S2 Algorithmic prediction of cryptic-activation. a-c)** DF (Donor Frequency), MES (MaxEntScan), NNS (NNSplice) and SAI (SpliceAI) scores for **(a)** *AM*-variants **(b)** *CM*-variants and **(c)** *AM/CM*-variants. Colour coding is explained in the Figure key. When a donor strength score of 0 is returned, we set it to 0.000001 to allow for the D calculations (VAR/REF; VAR = variant; REF = reference). **d)** Comparison of ***(i)*** NNS, ***(ii)*** MES and ***(iii)*** SAI scores with DF for all cryptic-donors (scores for VAR sequence) in our Cryptic-Donor database. DF shows strongest correlation with MaxEntScan. NNSplice does not recognise a subset of human donors to offer a strength prediction.

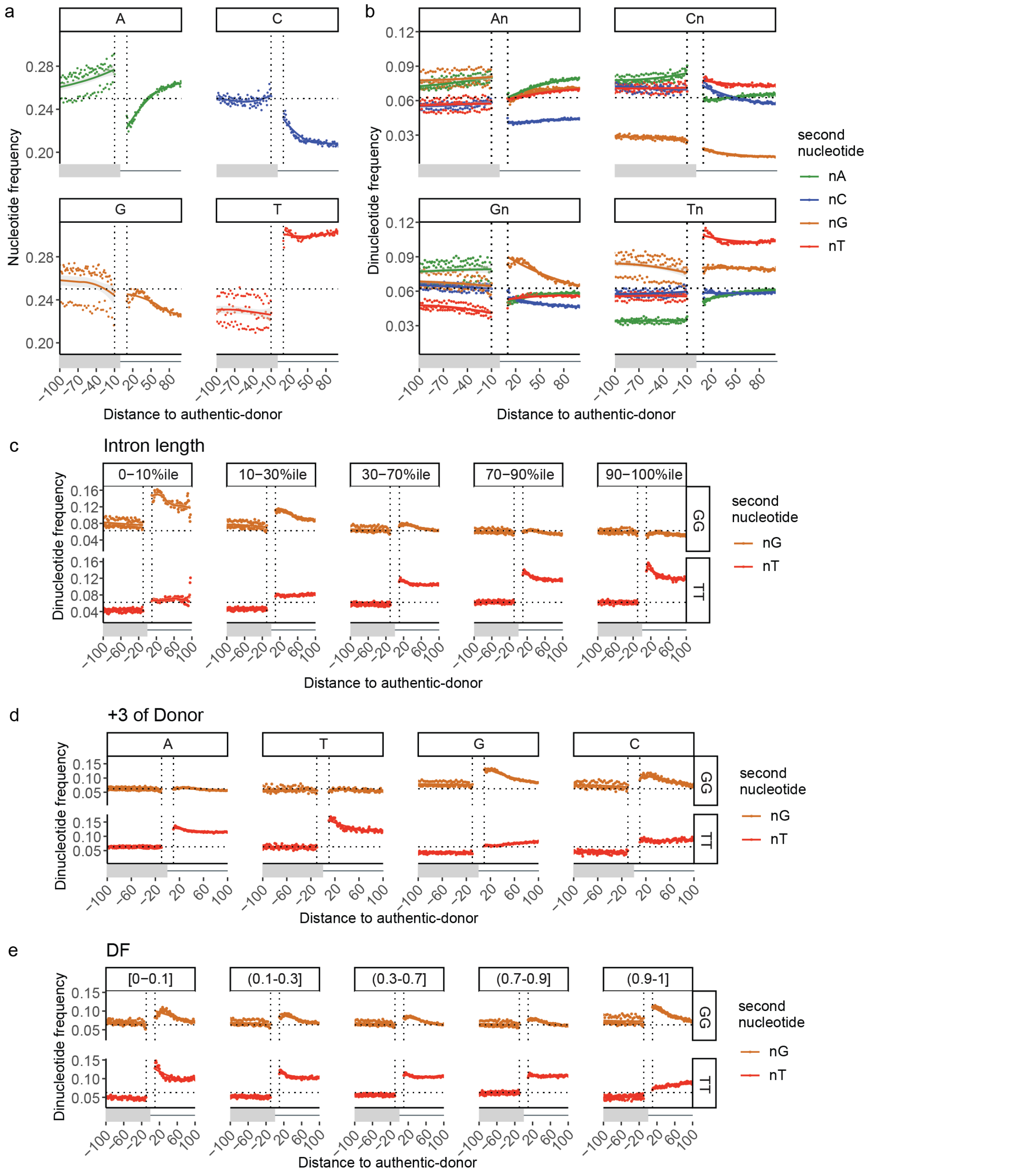


**Fig. S3 G- and T- dinucleotide repeats show distinct patterns of enrichment in different introns. a-b)** frequencies of each **(a)** nucleotide and **(b)** dinucleotide at each position surrounding authentic-donors. Vertical dotted lines denote boundaries at -10 and +10 where calculations start (i.e. excluding the conserved extended splice-site region). Horizontal dotted lines denote a random frequency of **a)** 1/4 for single nucleotides and **b)** 1/16 for dinucleotides. Lines show LOESS smoothing (locally weighted smoothing i.e. trendlines) with confidence bands in *grey*. In **(b)** Panels are according to the first nucleotide and colours are according to the second nucleotide in the dinucleotide. Note enrichment of G- and T- dinucleotides in the first 50 nt of the intron. **c-e)** frequencies of dinucleotides GG, and TT at each position surrounding authentic-donors. Vertical dotted lines denote boundaries at -10 and +10 where calculations start, horizontal dotted line denotes a random frequency of 1/16. Lines show LOESS smoothing (locally weighted smoothing i.e. trendlines) with *grey* confidence bands. **a)** G-repeats are enriched in the shortest human introns whereas T-repeats are enriched in longer introns. Length bins: < 149 nt (< $10^{th}$ percentile), 149-627 nt (10 - $30^{th}$ percentile), 628-3010 nt (30 - $70^{th}$ percentile), 3011-9270 nt (70 - $90^{th}$ percentile), > 9270 ntå (> $90^{th}$ percentile). **b)** Authentic donors with $D^{+3}$ G (or C) are enriched in G-dinucleotides whereas donors with $D^{+3}$ A (or T) are enriched in T-dinucleotides. **c)** Rare donors (low DF) show greater enrichment for T dinucleotide repeats compared with common donors.

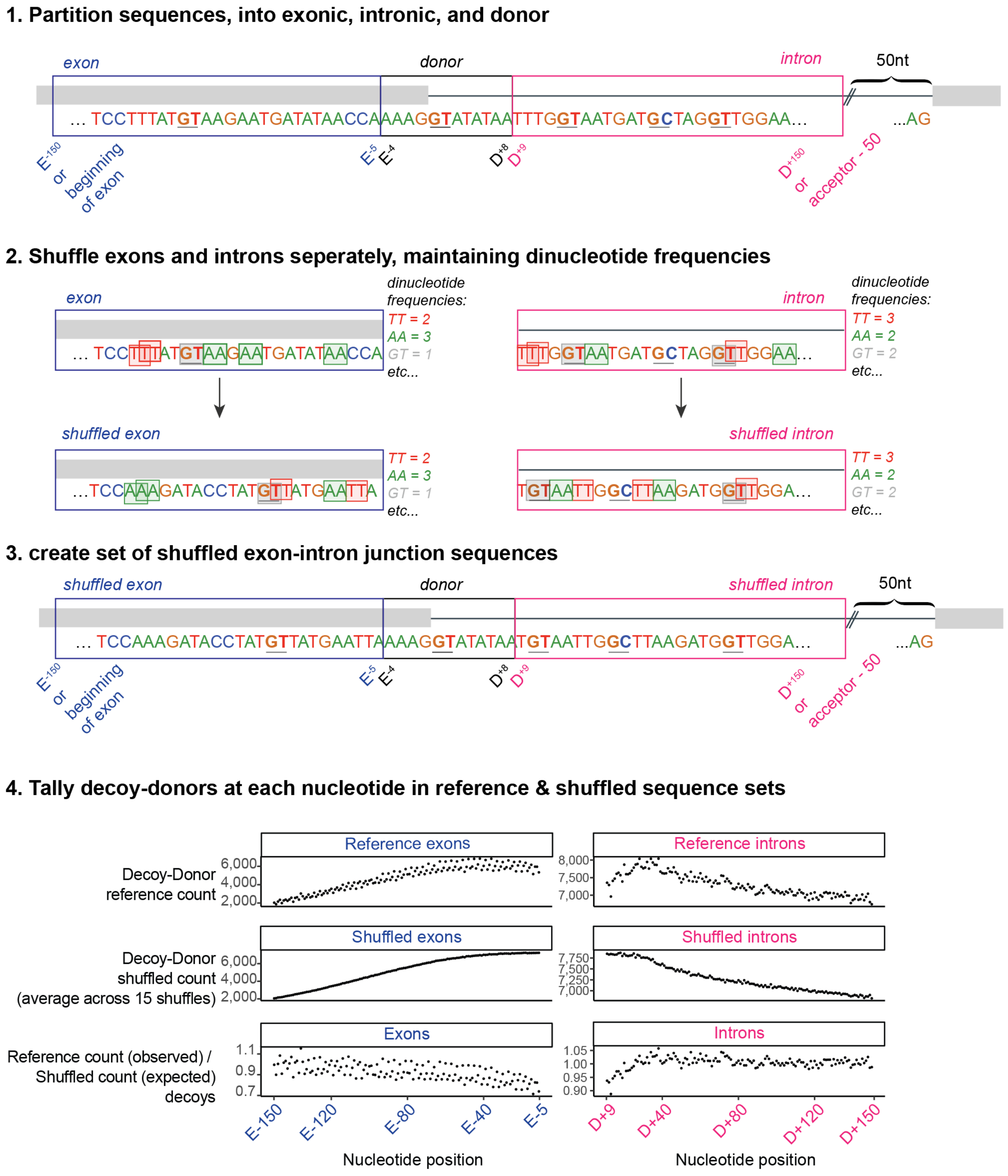


**Fig. S4 Schematic representation of how decoy depletion is calculated.** Sequences in *blue* 'exon' and *pink* 'intron' boxes are shuffled separately (maintaining dinucleotide frequencies) and the number of actual decoy-donors at each position is divided by the number in the shuffled sequence set.

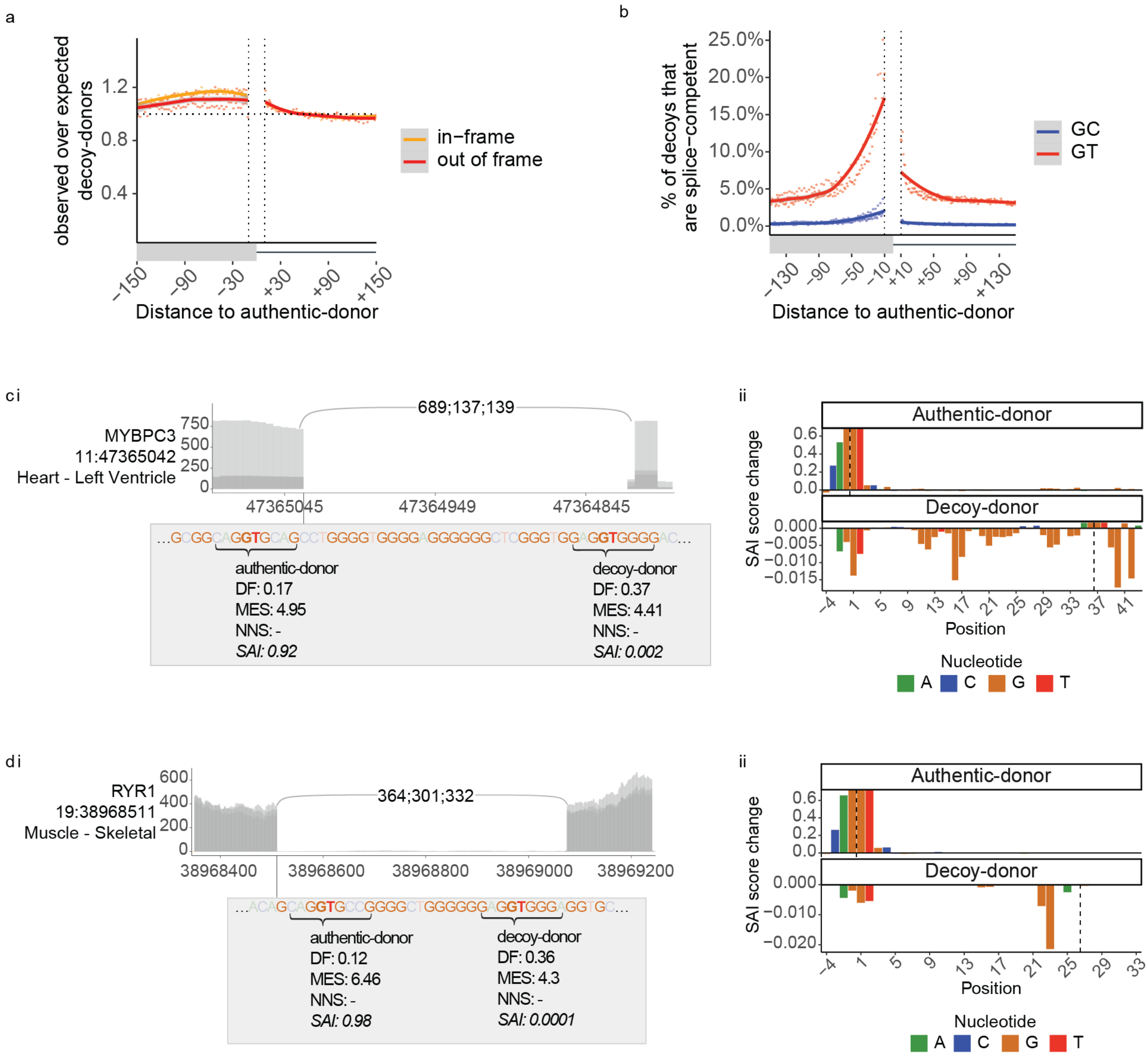


**Fig. S5 *Splice competence* provides insight into natural use of decoy-donors by the spliceosome. a)** The ratio of observed to expected 'GC' decoy-donors at each position +/-150 nt of authentic-donors in hg19. *Yellow*: In-frame exonic decoy-donors. *Red*: Exonic out-of-frame or intronic decoy-donors. **b)** The percent of decoys that are *splice competent* +/-150 nt of authentic-donors in hg19. *Red*: 'GT' decoy-donors show increasing *splice competence* with proximity to the authentic-donor. *Blue*: 'GC' decoy-donors show inherent low *splice competence*. At each position the number of *splice competent decoy-donors* (i.e. at least 1 read in aggregate RNA-Seq splice-junction data (see Methods) is divided by the total number of naturally occurring decoy-donors at that position. Lines show LOESS smoothing (locally weighted smoothing i.e. trendlines) with *grey* confidence bands. **c-d)** two examples of decoy-donors overlapping G-repeats that outcompete the authentic-donor according to DF and MES, but show no evidence for *splice competence,* and which SAI correctly identifies as non-functional donors. ***i)*** Shows overlays of 3 GTEx RNA-seq samples from the tissues with the highest TPM for that gene. The numbers (e.g. 689;137;139) denote the detected reads in each respective sample for that splice-junction. Algorithmic strength scores for authentic- and decoy-donors are boxed. ***ii)*** Result of SAI *in silico mutagenesis* showing the bases contributing to predicted strength of the authentic-donor (top) and decoy-donor (bottom). 'SAI score change' denotes the decrease (if positive) or increase (if negative) on the predicted strength of the donor when that nucleotide is mutated (see methods).

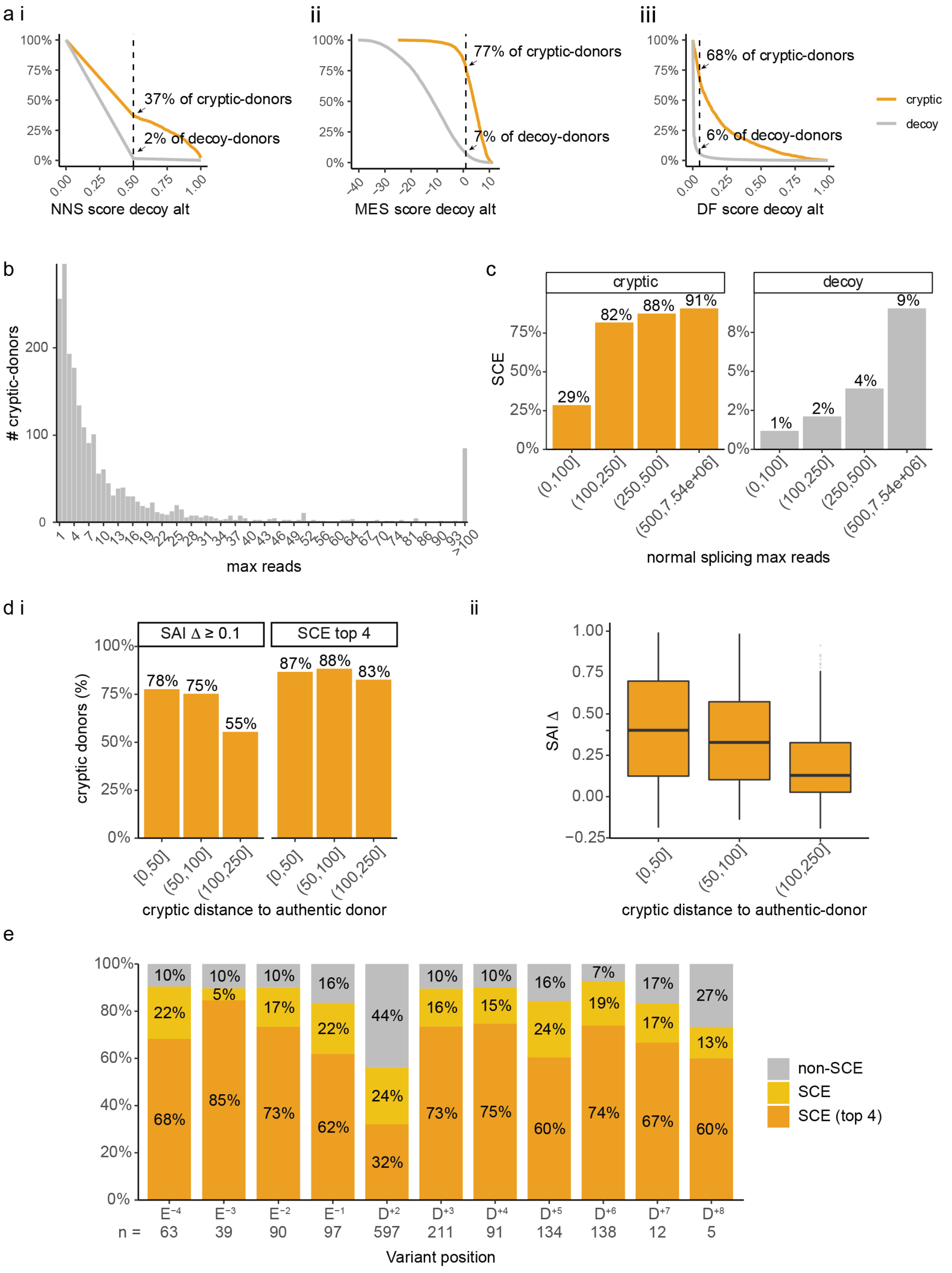


**Fig. S6 Metrics relevant to the employment of *splice competence* and SAI for prediction of cryptic selection. a)** Sensitivity (*orange*) and specificity (*grey*) of ***(i)*** NNS using a cut-off of 0.5, ***(ii)*** MES using a cut-off of 1 and ***(iii)*** DF using a cut-off of 0.05 to predict cryptic-donor activation in *AM*-variants. **b)** The maximum number of reads detected across 40,233 RNA-seq samples for each cryptic-donor activated by an *AM*-variant. **c)** Read-depth of the target gene influences sensitivity of SCE *(splice competent* events*)*. SCE predicts only 29% of cryptic-donors for target genes with < 100 max reads corresponding to normal splicing at the exon-exon junction under scrutiny, rising sharply to > 82% predictive accuracy with more than 100 max reads corresponding to normal splicing. **d*i)*** Percent of *AM-variant* cryptic-donors with SAI Δ scores greater than or equal to 0.1 (left) or in the SCE top 4 (right) in different bins according to cryptic distance to the authentic-donor. SpliceAI's ability to accurately identify cryptic-donors activated by *AM*-variants drops to 55% sensitivity for cryptic-donors more than 100 nt from the authentic-donor ***ii)*** SAI decoy-Δ scores for cryptic-donors relative to their distance from the authentic-donor. **e)** The percent of *CM*- and *AM/CM*-variant cryptics detected as *splice competent* events (SCE), according to the position of the SNV within the extended splice-site region of the activated cryptic-donor.
